# Soft skeletons transmit force with variable gearing

**DOI:** 10.1101/2024.03.28.587271

**Authors:** Olaf Ellers, Kai-Isaak Ellers, Amy S. Johnson, Theodora Po, Sina Heydari, Eva Kanso, Matthew J. McHenry

## Abstract

A hydrostatic skeleton allows a soft body to transmit muscular force via internal pressure. A human’s tongue, an octopus’ arm, and a nematode’s body illustrate the pervasive presence of hydrostatic skeletons among animals, which has inspired the design of soft engineered actuators. However, there is a need for a theoretical basis for understanding how hydrostatic skeletons apply mechanical work. We therefore model the shape change and mechanics of natural and engineered hydrostatic skeletons to determine their mechanical advantage (MA) and displacement advantage (DA). These models apply to a variety of biological structures, but we explicitly consider the tube feet of a sea star and the body segments of an earthworm, and contrast them with a hydraulic press and a McKibben actuator. A helical winding of stiff, elastic fibers around these soft actuators plays a critical role in their mechanics by maintaining a cylindrical shape, distributing forces throughout the structure, and storing elastic energy. In contrast to a single-joint lever system, soft hydrostats exhibit variable gearing with changes in MA generated by deformation in the skeleton. We found that this gearing is affected by the transmission efficiency of mechanical work (MA × DA) or, equivalently, the ratio of output to input work), which changes with the capacity to store elastic energy within helically wrapped fibers or associated musculature. This modeling offers a conceptual basis for understanding the relationship between the morphology of hydrostatic skeletons and their mechanical performance.

## INTRODUCTION

A diverse array of biological structures move by transmitting mechanical work through a soft body. This is achieved through hydrostatic skeletons, which are liquid-filled, pressurized structures that are actuated by layers of muscle. Examples include the human tongue, the body of worms and cnidarians, the mammalian penis (Chapman, 1958; Kier, 2012), and the body of a muscle, which itself generates intramuscular pressures that affect force transmission (Azizi et al., 2008; Kier, 2020; Sleboda and Roberts, 2020). Thus, hydrostatic skeletons are ubiquitous among animals and our understanding of their properties informs the field of biologically-inspired soft robotics (Kim et al., 2013; Rus and Tolley, 2015; Laschi et al., 2016; Hawkes et al., 2021). The study of hydrostatic skeletons has demonstrated how their geometry mediates the transmission of muscular displacement (Cowey, 1952; Clark and Cowey, 1958; Kier and Smith, 1985; Kier and Van Leeuwen, 1997; Van Leeuwen and Kier, 1997). However, it remains unclear how the transmission of force, and hence mechanical work, depends on the geometry and fiber winding of hydrostatic skeletons.

The gearing of a skeleton determines the extent to which muscular work is applied to displacement or force. Gearing is quantified by mechanical advantage MA, the ratio of output to input force, and displacement advantage DA, the ratio of output to input displacement. For a rigid skeleton, the MA about a joint may be shown from a balance of torques to equal the ratio of in-lever to out-lever lengths (Fig. 1F), which equals the inverse of DA (Smith and Savage, 1956). Neither metric changes with joint rotation and thus MA and DA are fixed properties of a skeleton’s geometry. As a consequence, measurements of MA and DA have routinely offered a basis for interpreting the functional morphology of vertebrate skeletons (Alexander, 1983; Biewener, 1989; Dunn, 2018; Westneat, 2003).

**Fig. 1.**
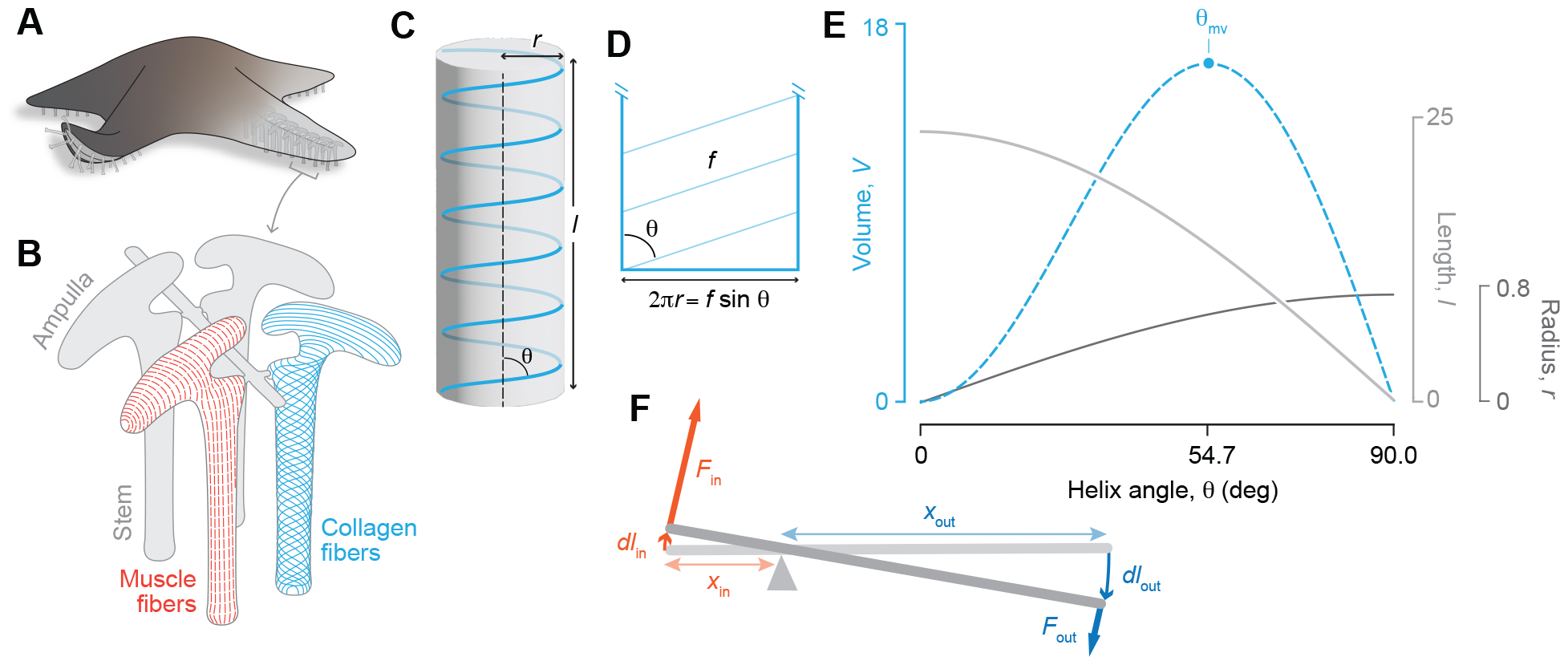
Skeletal force transmission depends on geometry. (A) The tube feet of sea stars are used for locomotion, adhering to substrata, feeding, and respiration. (B) Each tube foot is connected to an internal muscular bladder, the ampulla, and the tube foot-ampulla complex is wrapped in muscle (red) and collagen fibers (blue). The muscles are oriented in circular bands in the ampulla and in longitudinal bands in the tube foot; collagen fibers are oriented parallel to the long axis in the ampulla and cross-helically arranged in the stem (drawn based on McCurley and Kier, 1995). (C–E) The geometry of a helix informs models of hydrostatic skeletons that are reinforced with a cross-helical arrangement of collagen fibers. (C) The cylindrical volume (in gray) enclosed by a single helical fiber (in blue) illustrates how the helix angle (*θ*) relates to its radius (*r*) and length (*l*). (D) The surface of this volume shows the relationship between fiber length through one rotation (*f*), helix angle, and cylindrical circumference. (E) The cylindrical volume is plotted (dashed blue curve), along with its corresponding radius (dark gray, Eqn. 5) and length (light gray, Eqn. 4), as a function of helix angle for *n* = 5, and *f* = 1.5*π*, with maximum volume highlighted at *θ*_mv_. (F) A rigid lever system where the in-lever length (*x*_in_) is less than the out-lever length (*x*_out_) and hence where DA *>* 1 and MA *<* 1. Therefore, output force (*F*_out_) is less than input force (*F*_in_) and output displacement (*dl*_out_) is greater than input displacement (*dl*_in_).

Rigid skeletons comprised of multiple joints demonstrate how skeletal systems may change gearing as they deform. For example, the stomatopod raptorial appendage possesses a 4-bar linkage system that initially has a high MA that contributes to its rapid acceleration. As the strike advances, a reduction in MA enables input work to be imparted over a large displacement (McHenry et al., 2012). Flea legs similarly exhibit a high MA early in a jump as the body accelerates (Bennet-Clark and Lucey, 1967). In contrast, the initial low MA of a frog’s leg allows for storage of elastic energy prior to the principle body acceleration, but then increases over the jump for high displacement and work output (Astley and Roberts, 2014). Therefore, a change in gearing over the motion of an appendage has potential to enhance elastic energy storage, force output, and/or speed.

Geometric models of soft skeletons suggest that they also have variable gearing. Squid tentacles are a classic example of variable gearing. These tentacles capture prey by rapidly accelerating forward through contraction of circular and transverse muscles that pressurize the structure’s interior and reduce its radius. For a constant volume cylinder, length varies with the inverse square of radius. Consequently, the DA of a tentacle increases rapidly as it reaches full extension (Kier, 1982; Kier and Smith, 1985; Kier and Van Leeuwen, 1997). The extent to which variable gearing is a general feature of soft skeletons and whether MA is inversely related to DA, as previously proposed, remains to be determined (Kurth and Kier, 2014; 2015).

A complicating factor in the geometry and mechanics of soft skeletons is the common presence of a superficial layer of connective tissue. The connective tissue layer generally features a cross-helical arrangement of stiff collagen fibers that contribute to force transmission while providing reinforcement against aneurysms under pressure (Clark and Cowey, 1958; Wainwright, 1988; Shadwick, 2008). The helical arrangement factors into how these structures deform because a helix encloses a cylinder with a radius, length, and volume that varies with its pitch (Fig. 1C, D; Cowey, 1952; Clark and Cowey, 1958). The pitch, expressed by the helix angle *θ*, conforms to accommodate its interior volume.

The tube feet of echinoderms, used by sea stars for locomotion, offer an example of a variable volume hydrostatic skeleton with cross-helical fibers. A tube foot (i.e., podium; Smith, 1947; McCurley and Kier, 1995; Leddy and Johnson, 2000; Ellers et al., 2021) consists of a stem that extends from the body through pressurization of a lumen that is inflated by an internal muscular bladder called the ampulla (Fig. 1A–B). The stem and ampulla are each lined by a single muscle layer – circular in the ampulla and longitudinal in the stem – and the fluid exchange between the chambers allows these muscles to function as antagonists. This ability to generate extension depends on the helical winding around the stem. If initially retracted, a relatively high initial helix angle allows its value to reduce towards the value that maximizes the internal volume (54.7 deg, Fig. 1E). In support of this idea, the tube foot in one species of sea star has been shown to possess a helix angle of 67 deg at rest (McCurley and Kier, 1995). Engineers have exploited the converse case to develop a pneumatic muscle-inspired motor called a McKibben actuator (Chou and Hannaford, 1996; Liu and Rahn, 2003; Tondu, 2012). The flexible walls of a McKibben feature stiff fibers with a helix angle that begins at less than 54.7 deg, but increases during inflation to drive a contraction of the actuator’s length.

The present study develops an analytical framework for the transmission of mechanical work by soft skeletons. Our particular aims are to resolve whether variable gearing is common among a variety of hydrostatic systems and to determine the conditions under which MA is inversely related to DA. To this end, we model the geometry and mechanics in different hydrostatic skeletons to illustrate the effects of body deformation and helical winding on gearing. These models (summarized in Table 1) are of (1) an hydraulic press, (2) a McKibben actuator, (3) an echinoderm tube foot, (4) a cylindrical hydrostat without helical fibers, and (5) a cylindrical hydrostat with helical fibers. The first three of these are hydraulic systems where a fluid volume is exchanged between two chambers and the last two consist of a single chamber of fixed volume. The hydraulic press and McKibben actuator represent well-studied mechanical systems to which we can compare models of biological hydrostats.

**Table 1.** Summary of MA and DA for models.

| Model | MA (Eqn.) | DA (Eqn.) | MA × DA, $\eta$ |
| --- | --- | --- | --- |
| General | MA (2) | DA (1) | = 1 |
| Hydraulic press | MA<sub>press</sub> (10) | DA<sub>press</sub> (9) | = 1 |
| Piston-plus-McKibben | MA<sub>Mn</sub> (21) | DA<sub>Mn</sub> (12) | = 1 |
| Tube foot |  |  |  |
| No spring | MA<sub>tf</sub> (26) | DA<sub>tf</sub> (24) | = 1 |
| With spring | MA<sub>tf,spring</sub> (29) | DA<sub>tf</sub> (24) | < 1 |
| Cylindrical hydrostat |  |  |  |
| No fibers |  |  |  |
| Extension | MA<sub>E,cv</sub> (35) | DA<sub>E,cv</sub> (32) | = 1 |
| Retraction | MA<sub>R,cv</sub> (37) | DA<sub>R,cv</sub> (36) | = 1 |
| With fibers |  |  |  |
| Extension | MA<sub>E,cv,f</sub> (57) | DA<sub>E,cv,f</sub> (58) | < 1 |
| Retraction | MA<sub>R,cv,f</sub> (60) | DA<sub>R,cv,f</sub> (61) | < 1 |

### DA AND MA IN HYDROSTATIC SKELETONS

The geometry and mechanical properties of a skeleton determine how it will transmit input force and displacement. The skeleton is actuated by an input force (*F*_in_), that is exerted over some displacement (d*l*_in_) in the same direction to apply work (d*W*_in_) to the system. This work is transmitted through the skeleton to its output end, where force (*F*_out_) is applied over a displacement (d*l*_out_) in the same direction to generate output work (d*W*_out_). The amount of output displacement generated for a given input is determined by the DA, which is equal to the magnitude of the following derivative:

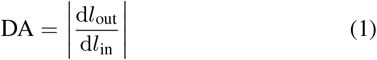

The choice of input and output directions is important in this calculation and should reflect the input and output work that are relevant to the situation. For instance, when an earthworm segment retracts, its input force will be generated by longitudinal muscles and its displacement would be the axial distance through which the muscles contract. Less obvious is the case of the input displacement for the segment’s circular muscles, where the input displacement is generated by a reduction in the circumference (i.e., d*l*_in_ = d*c* = 2*π*d*r*_in_). This contrasts with a reduction in the radial direction that may be directly generated by contraction of transverse muscles (i.e., d*l*_in_ = d*r*_in_) in some muscular hydrostats. The mechanical advantage also varies with the geometry of a system, but is defined as the ratio of the magnitudes of output and input forces:

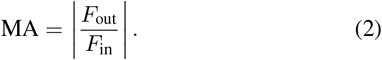

If no energy is stored or dissipated in the structure, then the output work and the input work are equal, and thus:

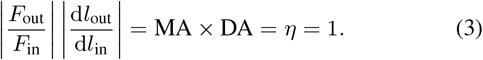

Therefore, the product of MA and DA provides a measure of the transmission efficiency *η* of mechanical work through a skeleton. A transmission efficiency of less than unity indicates either dissipation of mechanical energy from the system or storage in the form of elastic energy in the skeleton.

### THE GEOMETRY OF HELICAL FIBERS

Cross-helical stiff fibers transmit forces and play an important role in determining the mechanical and displacement advantage in a variety of soft skeletons. As in previous studies (Cowey, 1952; Clark and Cowey, 1958; Kier and Smith, 1985; Wainwright, 1988; Koehl et al., 2000; Kier, 2012), our models consider fibers wrapped helically around a cylinder at a helix angle *θ* from the longitudinal axis of the cylinder (Fig. 1C). In some cases below, we assume inextensible (i.e., fixed-length) fibers enclosing a variable volume (e.g., the McKibben and tube foot). In other models (e.g., the cylindrical hydrostat with helical winding), the volume is constant while the fiber length varies. In either case, we assume *f* is the length of a fiber through a single rotation of the helix (Fig. 1D) and *n* is the number of rotations of the helix on the cylinder. Then the cylinder enclosed by fibers at an angle *θ* has length *l*, radius *r*, and transverse cross-sectional area *A* given by:

**Table 2.** Symbols and subscripts.

| Symbols | Definition |
| --- | --- |
| $\epsilon$ | Strain |
| $\eta$ | Efficiency of energy transmission, $MA \times DA$ |
| $\theta$ | Helix angle |
| $\kappa$ | Ratio of wall volume to cylinder volume |
| $\sigma$ | Stress |
| $A$ | Area for force transmission |
| $E$ | Young's modulus |
| $a$ | Transverse cross-sectional area |
| $c$ | Circumference of cylinder |
| DA | Displacement advantage |
| $f$ | Length of fiber for single rotation of helix |
| $F$ | Force |
| $k$ | Spring stiffness |
| $l$ | Cylinder length |
| MA | Mechanical advantage |
| $n$ | Number of rotations of the helix on the cylinder |
| $P$ | Pressure |
| $V$ | Volume of fluid in a chamber |
| $\mathbb{V}$ | Volume of constant volume cylinder |
| $v$ | Volume of body wall |
| $r$ | Cylinder radius |
| $T$ | Tension |
| $t$ | Wall thickness |
| $W$ | Work |
| Subscripts |  |
| amp | ampulla |
| axial | along the longitudinal axis |
| circ | circular muscles |
| cv | constant volume cylindrical hydrostat |
| cvf | cv with helically wrapped fibers |
| E | extension |
| E=0 | cvf with fibers at zero stiffness |
| 54 | cvf with fibers at $\theta = 54.7$ deg |
| f | fiber |
| h | hoop |
| in | input |
| l | longitudinal |
| long | longitudinal muscles |
| m | muscle |
| Mn | piston-plus-McKibben |
| mv | maximum volume |
| o | equilibrium or resting value of a quantity |
| out | output |
| press | hydraulic press |
| Pi | piston of piston-plus-McKibben |
| R | retraction |
| spring | spring |
| tot | total |
| tf | tube foot |
| w | wall |

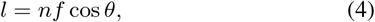

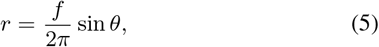

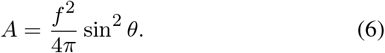

The enclosed cylinder’s volume using *V* = *πlr*^2^ with Eqns. 4 and 5 is:

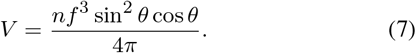

For the inextensible fiber case, plots of these variables (Fig. 1E) as shape changes, show the maximum volume occurring at 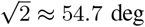. This fact was used in the first interpretation of the deformation of nemertean worms (Cowey, 1952) and generalized by Clark and Cowey (1958) for a broader diversity of worms.

The rate of change of length (Eqn. 4) with respect to volume (Eqn.7) is:

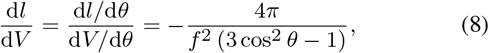

and reveals that length decreases if 0 *< θ < θ*_mv_ or increases if *θ*_mv_ *< θ < π/*2 as the volume approaches the maximum (Fig. 1E).

### THE HYDRAULIC PRESS

The properties of a hydraulic press are well-established (Miller et al., 2004), but we repeat them here for contrast with systems with deformable walls. The hydraulic press consists of rigid cylindrical input and output chambers filled with a constant volume of an incompressible fluid that is free to move between them. This fluid transmits forces from the input to the output pistons (Fig. 2A). The properties of the hydraulic press are determined by the geometry of thechambers. Due to their rigid walls, the transverse cross-sectional area of each chamber (*A*_in_, *A*_out_, respectively) is fixed and therefore the displacement of the two pistons (d*l*_in_, d*l*_out_) is related by the constant-volume condition. This yields the following displacement advantage (Eqn. 1):

**Fig. 2.**
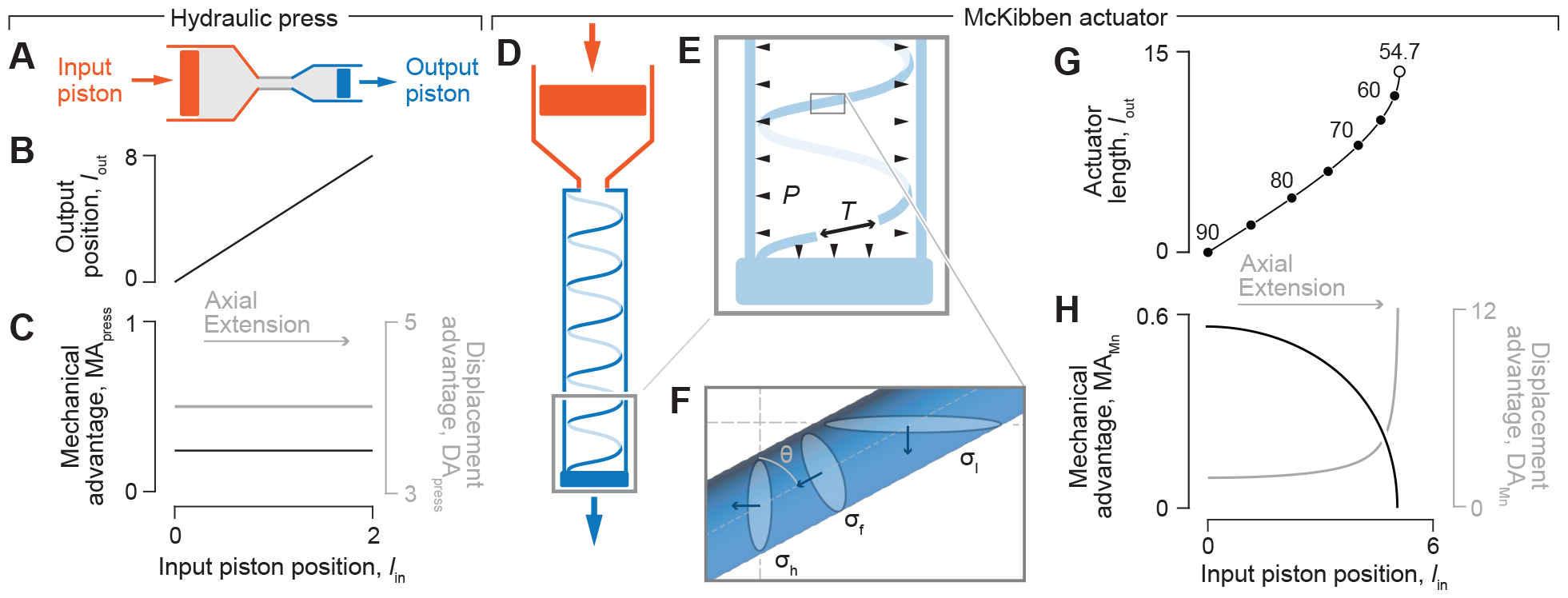
The transmission of force and displacement by a (A–C) hydraulic press and (D–H) McKibben actuator. (A) Schematic illustration of a hydraulic press. (B) The position of the output piston and resulting values for (C) MA_press_ (in black, Eqn. 10) and DA_press_ (in gray, Eqn. 9), as a function of the input position of a hydraulic press (Fig. 1A) where *A*_in_ = *π, A*_out_ = 0.25 *π*, and *P* = 1. (D) Schematic of the input chamber (in red) and fiber-reinforced hydraulic McKibben actuator (in blue) with a single helical fiber drawn. (E) An inset of the distal end of the McKibben actuator illustrates how pressure (*P*, arrowheads) acts on the inner walls and serves to generate the output force (blue arrow; Fig. 1D) and tension (*T*, black arrow) that resists extension. (F) An inset of a single fiber illustrates the relationship between fiber stress *σ*_f_, hoop stress *σ*_h_, and longitudinal stress *σ*_l_ with respect to helix angle *θ*. (G) Parametric plots of the McKibben actuator length (curve) and helix angle (in intervals of 5 deg, filled circles, and at 54.7 deg, open circle), as well as the (H) MA_Mn_ (in black) and DA_Mn_ (in gray) for *n* = 5, *f* = 1.5 *π*, and *A*_in_ = *π*, which were plotted as parametric functions of *θ* (Eqns 4, 12, 21, and 22). All parameter values are specified in arbitrary units.

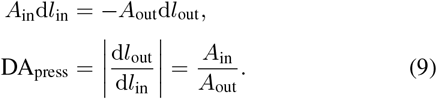

Assuming equal pressure *P* in the chambers, the mechanical advantage (Eqn. 2) for the hydraulic press is:

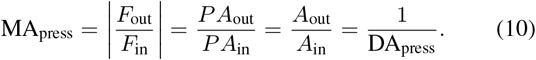

Thus, transmission efficiency for the hydraulic press is:

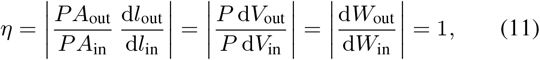

where *W*_out_ and *W*_in_ are work-out and work-in, respectively. The inverse relationship between MA_press_ and DA_press_ is a manifestation of the complete transmission of energy from the input to the output chambers. In addition, the displacement of the output piston is directly proportional to that of the input piston (Fig. 2A) and hence MA_press_ (Eqn. 10) and DA_press_ (Eqn. 9) are fixed across all positional changes (Fig. 2C), as in a lever (Fig. 1F).

### THE MCKIBBEN ACTUATOR

The McKibben actuator consists of a deformable fluid chamber constrained by inextensible, cross-helical fibers (Fig. 2D). We model the deformable McKibben as actuated by the piston of a rigid input chamber with both chambers filled by an incompressible fluid. Assume that the volume leaving the piston chamber is the same as the volume entering the McKibben chamber and use

Eqns. 1 and 8 to calculate DA_Mn_:

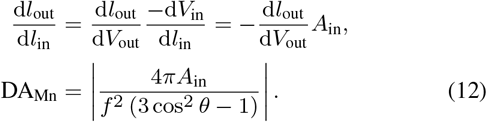

In contrast to the hydraulic press, where DA_press_ is constant as the piston pushes (Eqn. 9), DA_Mn_ changes as the McKibben actuator changes volume and length, which alters gearing. For an elongating McKibben actuator (i.e., *θ*_mv_ *< θ <* 90 deg), length (Eqn. 4) increases until the helix accommodates the maximum volume at *θ* = *θ*_mv_. Thus, DA_Mn_ (Eqn. 12) increases rapidly near to that maximum-volume full length (Fig. 2H).

To determine mechanical advantage of the piston-plus-McKibben MA_Mn_, we first calculate axial output force. We derive this force, previously derived using an energy argument (Chou and Hannaford, 1996), by considering stress in the helical fibers. We assume inextensible fibers that carry all forces in tension, with no contribution from any other material in the cylinder wall. We relate stress in the helical fibers *σ*_f_ to hoop stress *σ*_h_ and longitudinal stress *σ*_l_ in the cylinder wall (Fig. 2F):

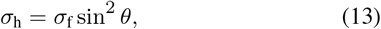

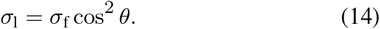

Consider a force balance across a longitudinal plane of the cylinder with wall thickness *t*:

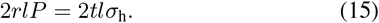

Using Eqn. 13 in Eqn. 15, stress in the helical fibers is:

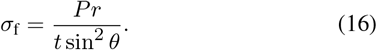

The longitudinal output force is the sum of the force due to pressure on the end of the cylinder minus the horizontal component of the force exerted from the helical fibers in the wall on the cylinder ends:

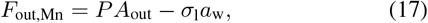

with area of the transverse section of the cylinder wall being:

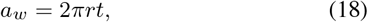

which assumes a thin wall as in Demirkoparan and Pence (2015). The output force for the piston-plus-McKibben system, using Eqns 5, 6, 14, 16 and 18 with Eqn. 17. is:

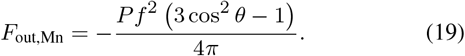

This relationship shows that zero force is generated at *θ*_mv_, negative (i.e., pulling) forces are generated where 0 deg *< θ < θ*_mv_, and positive (i.e., pushing) forces occur where *θ*_mv_ *< θ <* 90 deg. Note that our sign convention for forces is the opposite of Chou and Hannaford (1996).

The input force on the piston and the internal pressure in the machine are related:

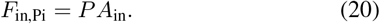

Mechanical advantage for the piston-plus-McKibben compound machine (using Eqn. 19) is thus:

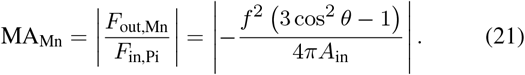

Therefore, by comparison to its displacement advantage (Eqn. 12), MA_Mn_ = 1*/*DA_Mn_ and mechanical advantage hence declines towards zero as displacement advantage increases near the full extension of the actuator. This inverse relationship indicates an ideal transmission efficiency (*η* = 1) and complete conversion of input work to output work by the actuator.

We used parametric plots to illustrate how the piston-plus-McKibben compound machine exhibits variable gearing as it is inflated (Figs. 2G–H). We evaluated DA_Mn_, MA_Mn_, and *l*_out_ (Eqns 12, 21, and 4, respectively) over the range of helix angles that correspond to elongation of the actuator (90 deg*> θ > θ*_mv_). These quantities are plotted against displacement of the piston (*l*_in_) as a function ofhelix angle using the assumptions of constant-volume, fluid incompressibility, and rigid input-chamber geometry:

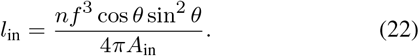

These plots illustrate that the actuator length increases more rapidly with input displacement at greater extension (Figs. 2G). This trendresults in an increase in DA_Mn_ and a corresponding decrease in MA_Mn_ as the actuator extends (Figs. 2H). Therefore, the McKibben actuator is most effective at transmitting force at low levels of inflation and is geared more for displacement when inflated close to its maximum volume.

### THE TUBE FOOT

We model the tube foot as a two-chambered system consisting of ampulla and stem. The stem of the tube foot is a fiber-wound cylindrical structure (Fig. 1B, 3A) that we model as a McKibben actuator (Figs. 2D–H). We model the compliant ampulla as a cylinder with a fixed length (*l*_amp_) and variable circumference (*c*_amp_), consistent with previous work (Kier, 2012). The compliant ampulla distinguishes the tube foot from the piston-plus-McKibben. However, as for the McKibben, we relate volumetric changes of input and output chambers. Specifically, assuming a constant volume sum of ampulla (here *V*_in_) and tube foot (*V*_out_, Eqn. 7), the circumference of the ampulla, as a function of helix angle is:

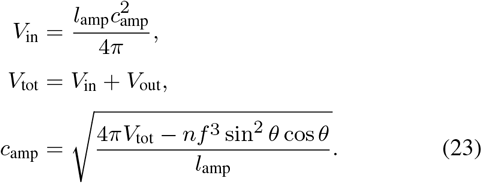

Circumference is the relevant dimension of the input chamber when calculating displacement advantage because work is generated by contraction of circumferential muscles. Displacement advantage (Eqn. 1) is the rate of stem extension with respect to changes in the ampulla’s circumference (Eqns 8 and 23):

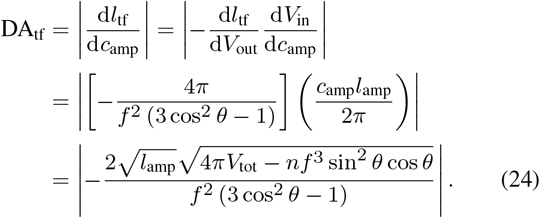

The compliant ampulla additionally plays a role in the mechanical advantage. Circular muscles of the ampulla generate an input force equal to the product of tension *T* and the length of the ampulla (*F*_in_ = *Tl*_amp_). Using Laplace’s Law for hoop tension in a cylinder, one may relate this muscular tension to the circumference of the ampulla (*P* = 2*πT/c*_amp_). Therefore, pressure generated by the ampulla is:

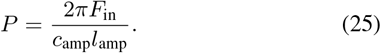

Next, we determine the mechanical advantage in two cases - with and without energy storage by elastic elements in the structure.

### Without elastic energy storage

Assume that all input work is transmitted to the output work. Combining our models for the output force (Eqn. 19) and pressure (Eqn. 25), gives the mechanical advantage for the tube foot MA_tf_:

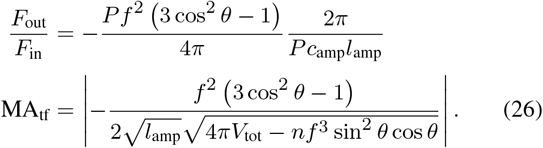

Therefore, MA_tf_ = 1*/*DA_tf_, as expected for a system without energy loss or storage. As a tube foot extends, helix angle will decrease and a maximum extension will occur at *θ* = 54.7 deg, at which point mechanical advantage becomes zero and no output force is generated. Longer tube foot lengths will only occur if the tube foot sucker is pulled by an external force to lengthen the tube foot further.

DA_tf_, MA_tf_, and *l*_tf_ (Eqns 24, 26, and 4, respectively) may be plotted parametrically with respect to *c*_amp_ (Eqn. 23) over the range of helix angles that correspond to elongation (*θ*_mv_ *< θ <* 90 deg, Fig. 3). As circumference decreases, the tube foot lengthens and helix angle decreases (Fig. 3B). As in the piston-plus-McKibben compound machine, these patterns result in an elevation of DA_tf_ and reduction of MA_tf_ near full extension of the tube foot (Fig. 3C). However, the gearing of the piston-plus-McKibben shows a monotonic decrease in mechanical advantage (Fig. 2H) as it lengthens, whereas the tube foot has a maximum mechanical advantage that occurs at some intermediate length as the stem extends (Fig. 3C). That maximum occurs at a circumference that depends on the relative size of the ampulla and tube foot stem.

**Fig. 3.**
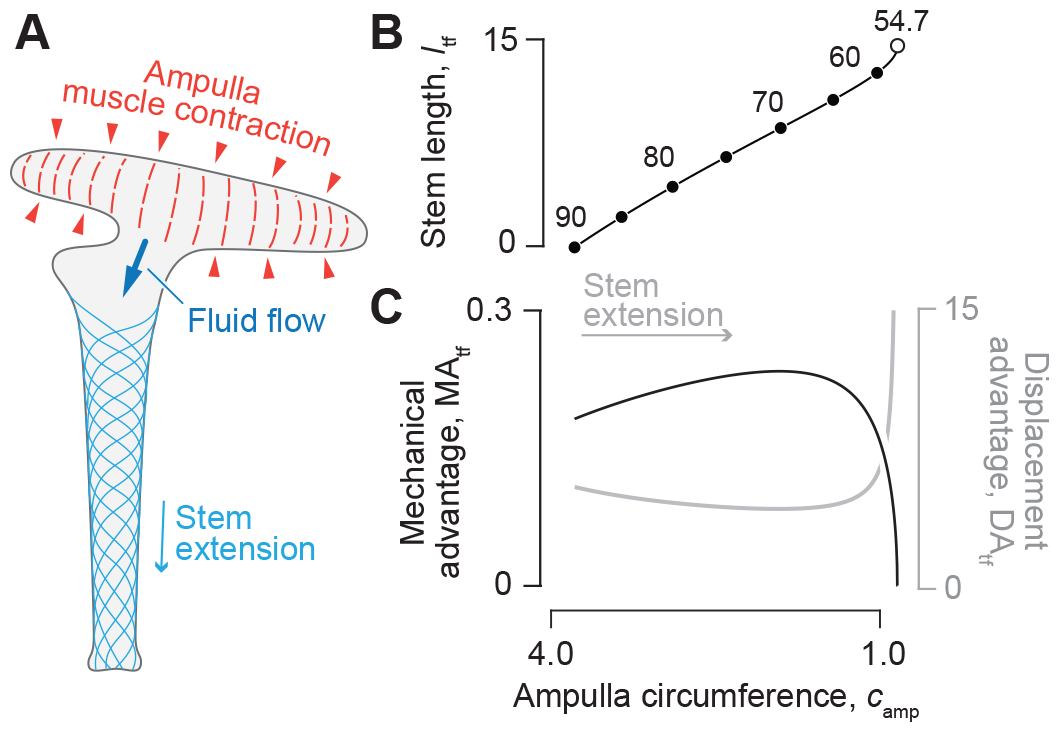
Force and displacement transmission in the tube feet of sea stars. (A) The tube foot extends through contraction of ampullar muscles, which pressurizes the chamber and induces flow into the tube foot stem. Parametric plots illustrate the change in (B) tube foot length and fiber helix angle (intervals of 5 deg, filled circles, and at 54.7 deg, open circle) during extension for *n* = 8, *l*_amp_ = 7, *f* = *π*, and *V*_tot_ = 8. The associated changes in (C) MA_tf_ (black) and DA_tf_ (gray) are shown with respect to that reduction in ampullar circumference *c*_amp_, which drives tube foot extension. All curves were plotted as parametric functions of *θ* (Eqns 4, 24, and 26) versus *c*_amp_ (Eqn. 23), all in arbitrary units.

### With elastic energy storage

Here, we consider the effects of energy storage due to passive force from longitudinal muscles in the stem (Fig. 1B). We speculate that the longitudinal muscles are initially folded when the stem is retracted and after reaching the muscles’ unfolded length, start generating a passive linearly elastic tensile force in proportion to its length beyond that point. This force counters the extension of the tube foot beyond the unfolded length, but offers no resistance at shorter lengths. This spring-like force is:

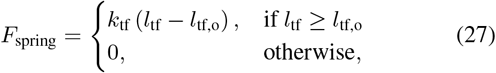

where *k*_tf_ is the stiffness of the tube foot spring (Fig. 4A). This spring approximates the passive elastic component of the longitudinal muscles and not their active tension. The spring resting angle *θ*_out,o_, as a function of *l*_tf,o_, can be calculated from the length of the helix (Eqn. 4):

**Fig. 4.**
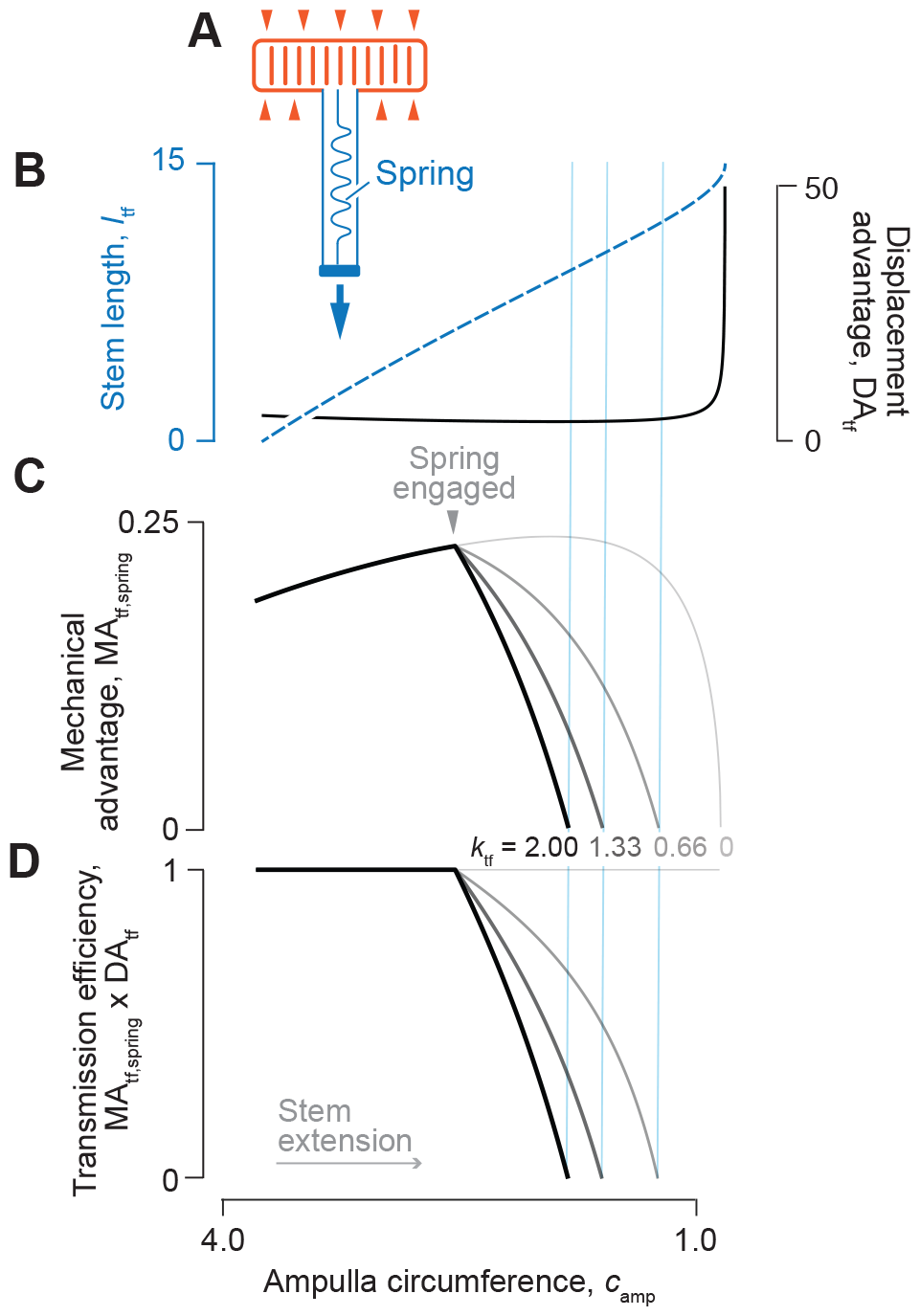
The effects of a parallel elastic element on transmission of force and displacement in the tube foot. (A) A schematic drawing of a tube foot that is extending due to the action of force (red arrows) generated by circular muscles, with a parallel elastic element (i.e., spring) that runs the length of the tube foot. (B) The change in tube foot length (dashed blue) and DA_tf_ (solid gray) are identical to what is depicted in Fig. 3 due to the calculations using the same parameter values (*n* = 8, *l*_amp_ = 7, *f* = *π*, and *V*_tot_ = 8). However, values for (C) MA_tf,spring_ vary with the stiffness and resting length of the tube foot spring, with stiffness of each curve between *k*_tf_ = 0 (lightest gray) to *k*_tf_ = 2 (black), where *l*_tf,o_ = 6. The spring is engaged in the middle of extension (gray triangle) and the vertical blue lines denote the values of the ampulla circumference where MA_tf,spring_ = 0, which represents a limit to the possible stem lengths and values of DA_tf_ that may be attained in (B). (D) The corresponding product of MA_tf,spring_ and DA_tf_ is shown for each spring stiffness value, where deviation from unity (i.e. *η <* 1) illustrates conditions where MA_tf,spring_ ≠ 1/DA_spring_. All variables were calculated as parametric functions of *θ* (Eqns 4, 24, and 29), with parameter values specified in arbitrary units.

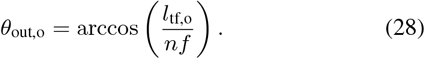

Accounting for force exerted by this spring (Eqn. 27) in (Eqn. 26) yields a modified mechanical advantage:

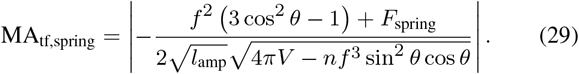

This equation for the mechanical advantage deviates from the inverse of DA_tf_ (Eqn. 24) for any non-zero spring stiffness, when the tube foot extends beyond the resting length of the spring.

This is because the passive elasticity of the longitudinal muscle stores some of the energy generated by the ampulla, rather than allowing all work to be transmitted to extending the tube foot. Once the ampulla relaxes, that stored elastic energy may then contribute to passive shortening of the tube foot. However, if the resistance to lengthening were instead a consequence of damping by the longitudinal muscle (not presently modeled), then the energy dissipation would reduce the output force but not contribute to subsequent retraction.

We illustrate the properties of the tube foot with the internal spring using parametric plots of the relevant equations over the same range of helix angles as previously. While stem length (*l*_tf_) and displacement advantage (DA_tf_) are independent of the effect of muscle spring constant (Fig. 4B), mechanical advantage (MA_tf,spring_) changes with the spring constant (Fig. 4C; Eqn. 29). Consequently, once the spring engages, transmission efficiency (i.e., MA_tf,spring_ × DA_tf_; Fig. 4D) varies as the stem extends. This is an example where energy storage in an elastic element results in *η <* 1.

The addition of an effective spring reduces the maximum elongation of the tube foot achievable through ampullar contraction, presenting an intriguing mechanism to control output displacement of McKibben-type systems. As a McKibben actuator without an internal longitudinal spring approaches its maximum volume, displacement advantage diverges rapidly, meaning that even a small change in input length will correspond to a large change in output displacement. A sufficiently stiff spring could reduce the operating length of the McKibben actuator such that displacement advantage is nearly linear over the full range, which may allow more precise control of output displacement.

### THE FIBERLESS CYLINDRICAL HYDROSTAT

The cylindrical hydrostat includes a constant volume of incompressible fluid or tissue volume lined with longitudinal and circular muscles. Fiberless models have been previously employed to consider the displacement advantage of a squid’s tentacle (Kier, 1982; Kier and Smith, 1985; Kier and Van Leeuwen, 1997) and the fluid-filled body of an earthworm (Kurth and Kier, 2014; 2015). In these models, either transverse and circular muscles (e.g., the squid) or circular muscles alone (e.g., the earthworm) cause axial extension of the body (Fig. 5A). Any differences in the gearing between circular and transverse muscles would help to indicate the functional implications of their presence and the circumstances where it may be beneficial for their activation on extension. The longitudinal muscles reverse this extension and apply a radial force by pressurizing the body’s interior and causing radial expansion. This radial force may act on the surrounding environment (Fig. 5D), which permits an earthworm to anchor some segments against a burrow (Quillin, 2000). Here, we neglect any internal resistance from stretching skin and hence the radially-directed forces act entirely on the environment. As we demonstrate, the mechanical advantage and displacement advantage depend on whether circular or transverse muscles are generating the extension.

**Fig. 5.**
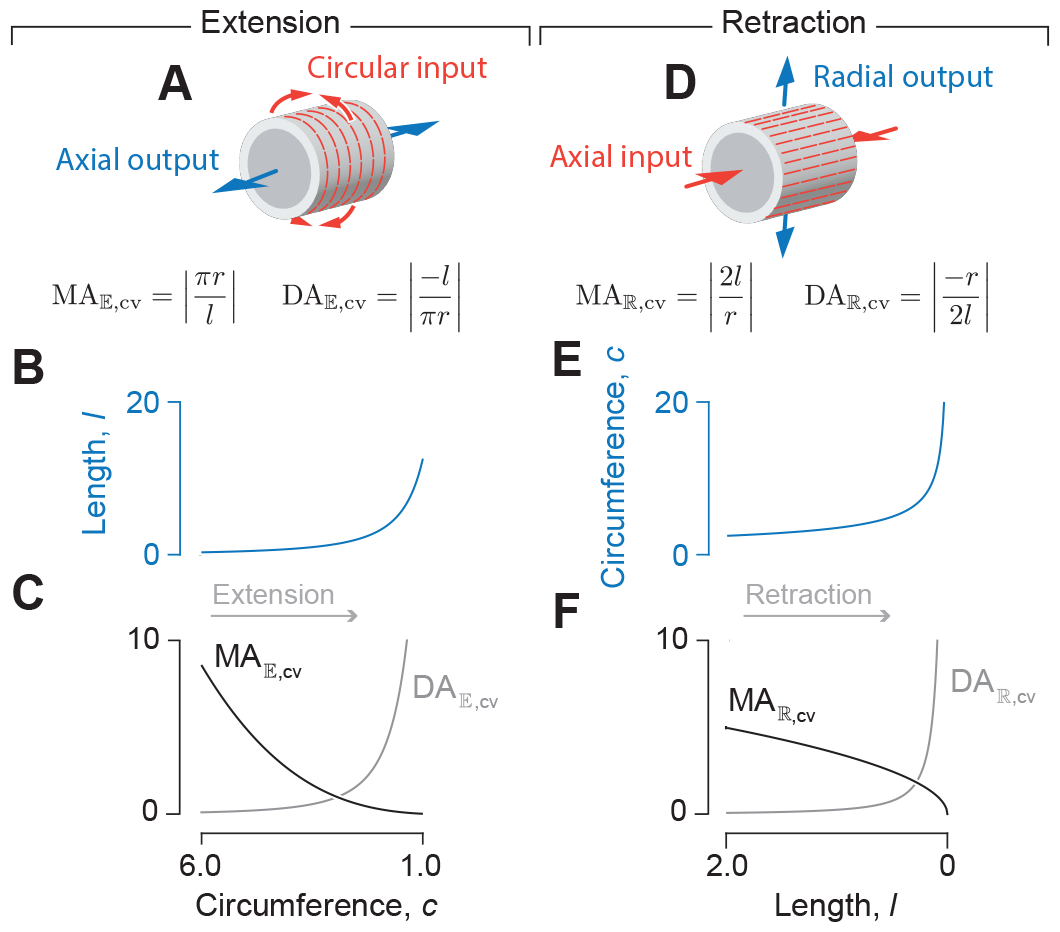
Transmission of force and displacement by a cylindrical hydrostat of constant volume. (A–C) Changes in dimensions and gearing on axial extension. (A) Schematic illustration of contraction by the circular muscle (input force, in red), which generates axially-directed output force and displacement (blue arrows). The change in (B) length (Eqn. 30), (C) MA_𝔼,cv_ (black, Eqn. 35), and DA_𝔼,cv_ (gray, Eqn. 32) varies with circumference. (D–F) The same variables are shown for axial extension. (D) Schematic illustration of the radial output force and displacement (blue arrows) generated by a contraction of longitudinal muscles (axial input, in red). (E) Circumference (Eqn. 31) of the hydrostat increases with decreasing length, which drives (F) decreases in MA_ℝ,cv_ (black, Eqn. 37) and increases in DA_ℝ,cv_ (gray, Eqn. 36). All parameter values are specified in arbitrary units, with 𝕍 = 1 and *n* = 1.

The displacement advantage of a cylindrical hydrostat derives from its length *l* and radius *r*:

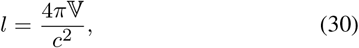

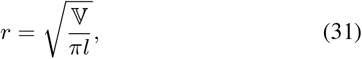

where 𝕍 and *c* are, respectively, the volume and circumference of the constant volume cylinder. The displacement advantage for the extension of a cylindrical body by the contraction of circular muscles is the derivative of length with respect to circumference:

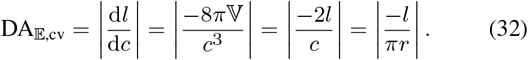

This length to radius ratio increases rapidly as the body becomes more elongated (Fig. 5B,C). If the muscles doing work were transverse, then the relevant input length would be diameter *d* instead of circumference, in which case we would take the derivative of length with respect to diameter which gives a displacement advantage that is a factor of d*c/*d*d* = *π* larger than that in Eqn. 32. Thus, the transverse muscles of a squid tentacle offer substantially greater amplification of displacement during extension, and correspondingly less mechanical advantage, than would circular muscles.

We now consider the mechanical advantage for extension based on the contraction of circular muscles. This force is generated by hoop tension, *T*_h_, which is related to the pressure in the cylinder *P* by Laplace’s Law:

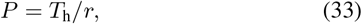

Thus the input force is:

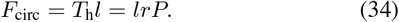

The axial force output at the cylinder’s end is the product of pressure and circular cross-sectional area of the cylinder; therefore, the mechanical advantage is:

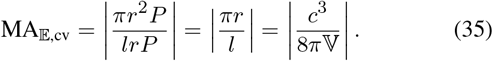

Thus, MA_𝔼,cv_ = 1*/*DA_𝔼,cv_ (Eqn. 32), as must be the case in the absence of energy dissipation or storage. If the muscles generating the input force were transverse instead of circular, then the mechanical advantage would be lower than Eqn. 35 by a factor of 1*/π*.

During retraction, activation of the longitudinal muscles serves to shorten the body and thereby increase its radius. Given the relationship of the radius to the volume of the body (Eqn. 31), the displacement advantage i:

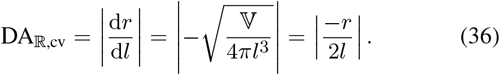

Therefore, the displacement advantage increases rapidly as the longitudinal muscles contract (Fig. 5E–F).

The mechanical advantage for retraction considers the radially-directed output force that is transmitted by pressure. The input force generating the pressure is created by the longitudinal muscles acting over the cylinder ends. The mechanical advantage in retraction is thus:

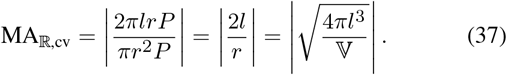

Therefore, MA_ℝ,cv_ = 1*/*DA_ℝ,cv_ (Eqn. 36) and MA_ℝ,cv_ declines during retraction (Fig. 5F).

When comparing mechanical and displacement advantage between extension and retraction, it is crucial to identify muscle orientation, relevant input and output directions and distances over which work is done (Fig. 5).

### THE CYLINDRICAL HYDROSTAT WITH ELASTIC CROSS-HELICAL FIBERS

Cylindrical hydrostats generally possess a cross-helical winding composed of collagen fibers (Clark and Cowey, 1958; Wainwright, 1988; Shadwick, 2008). The helix angle of these fibers is approximately 54.7 deg in the body wall of a diversity of worms (Clark and Cowey, 1958; Kier, 2012), some polychaetes (Law et al., 2014), and in the notochords of developing vertebrates (Adams et al., 1990; Koehl et al., 2000). As for hydraulic systems, one mechanical role of helical fibers in hydrostats is thought to aid in maintaining its cylindrical shape as it deforms (Wainwright, 1988). However, forces transmitted by these fibers have not previously been explicitly included in models of helically wound cylindrical hydrostats. Here, we consider the geometry and mechanics of fiber winding around a constant-volume cylindrical hydrostat and we develop expressions for mechanical advantage, displacement advantage, and transmission efficiency.

### Helical geometry with changes in fiber length

A pressurized helically wrapped, constant volume cylinder will have the fibers at an angle of 54.7 deg in the absence of external forces, and a change in shape will tend to reduce the volume (Fig. 1E). Therefore, for an incompressible object where volume cannot change, fibers must extend. For example, the body segment of an earthworm that is filled with a volume of fluid accommodated by a helix angle of 54.7 deg requires that the fiber change length for the segment to change in length.

The fiber length per turn of the helix is found by rearranging the equation for the volume enclosed by the helix (Eqn. 7):

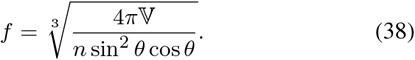

The resting fiber length per rotation of the helix (*f*_o_) occurs at the equilibrium angle (*θ*_mv_ = 54.7 deg) when there are no external longitudinal or radial forces, giving:

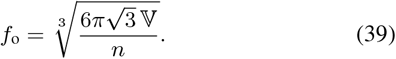

The change in fiber length from this resting length to achieve a particular helix angle is the difference between the fiber length for that angle (Eqn. 38) and the resting fiber length (Eqn. 39). Given variable fiber length and constant volume, the cylinder’s length, radius, and cross-sectional area with respect to helix angle are calculated by substituting Eqn. 38 into Eqns 4, 5, 6:

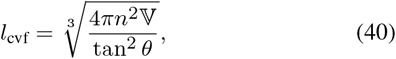

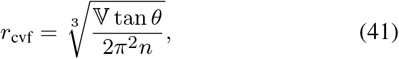

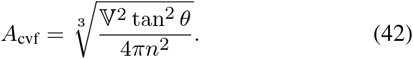

Note that, in all derivations, we assume that physical parameters are greater than zero and that fiber angle is 0 *< θ < π/*2.

Next, we calculate the ratio of length to radius in the context of a fiber angle. This variable was important in calculating the mechanical and displacement advantage for cylindrical hydrostats without fibers, and we now want to be able to compare hydrostats with and without fibers. Using Eqns. 40 and 41:

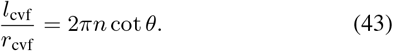

Thus the shape of the cylinder depends on both the number of wrappings and the fiber angle. Consider the geometry for earthworms. Average length to diameter ratio for whole earthworms was 102*/*5.3 and they had an average of 145 segments (Quillin, 1998) yielding a length to radius ratio of 0.27. This ratio at a fiber angle of 54.7 deg in Eqn. 43 implies that *n* = 0.06; i.e. the helical fiber wraps each segment just 0.06 times. That a segment is wrapped only 0.06 times by a helical fiber at a 54.7 deg angle is a surprising and testable prediction of the assumed geometry. A longer structure such as a squid tentacle with length to radius ratio of about 40 at that same angle (Kier and Smith, 1985) would be wrapped 9 times.

### Assume a constant volume of body wall

The body wall deforms as a cylindrical hydrostat changes shape. To account for this deformation, we assume that the body wall is constant volume and thin relative to the radius of the cylindrical body. The thickness of the body wall is therefore given by:

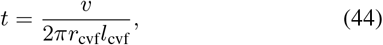

with *v* being the volume of the body wall. This body wall has a cross-sectional area:

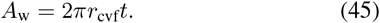

We assume that the volume of the body wall is a fraction *κ* of the body volume:

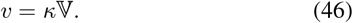

Under these assumptions, the ratio of the body wall thickness (Eqn. 44) to the radius (Eqn. 46) is:

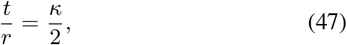

which is a constant for all worm shapes. In our example calculations, we use *κ* = 0.1 which means that the wall thickness is 0.05 times the radius.

### The mechanics of linearly elastic fibers

Fiber extension depends on the material properties of the fibers. We assume a constant stiffness and hence a model of fibers that generate force in proportion to their extension beyond a resting length. Therefore, stress in a fiber with strain *ϵ* and Young’s modulus *E* is:

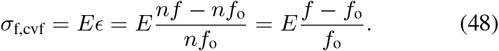

By substitution of fiber length (Eqns. 38 and 39), the stress, with respect to the helix angle, is:

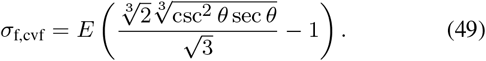

For this expression, the components of stress along the hoop and longitudinal directions (Eqns 13 and 14) are:

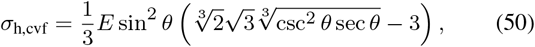

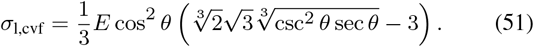

We can relate the fluid pressure to the fiber stress. This is achieved through considering a balance of forces across a longitudinal plane through the worm, similar to the McKibben (Eqn. 15), but including the hoop stresses *σ*_m_ generated by circular muscles:

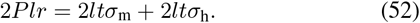

The muscles here are assumed to produce a fixed level of stress, without consideration of the muscle’s length-tension curves or force-velocity properties. Because muscles can control their level of activation, a muscle can control the stress produced, to some extent, independent of the point on the length-tension curve at which the muscle is operating. We are primarily interested in the way that a hoop fiber or muscle can carry some of the hoop stress, and we are interested in how that changes the angle at which forces are balanced. Therefore we model only this one aspect of circular muscle action - the level of stress. This allows us to generate graphs that are series of static balances with different fiber angles and output and input forces. The model we build can be subsequently modified to model muscle properties more specifically.

From this force balance across the longitudinal plane (Eqn. 52), we incorporate our expressions for the hoop stress of fibers (Eqn. 50) and the body dimensions (Eqns 40, 41, and 44):

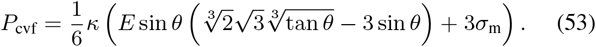

This relationship between pressure and body shape reveals some fundamental mechanical features. In the absence of circular muscle stress (*σ*_m_ = 0), the internal pressure is minimized when the body is at a circumference where *θ* = 54.7 deg (Fig. 6D). Pressure is higher at fiber angles on either side of this minimum pressure angle. Thus, a worm that is extending or retracting will generate internal coelomic pressures consistent with experiments (Seymour, 1969; Quillin, 1998; 2000). Circular muscle contraction with concurrent elongation coincides with increases in coelomic pressure, whereas subsequent circular muscle relaxation and contraction of the longitudinal muscle cause a decrease in pressure as the angle returns to 54.7 deg.

#### The axial force with cross-helical fibers

The axial force is the input to the hydrostatic skeleton on retraction and the output on extension. This force could be due to a combination of longitudinal muscles, an external force, or stress within the fiber winding. We found an equation for the axial force by considering internal pressure and resistance generated by the fiber winding, which mirrors our derivation of output force by the McKibben (Eqn. 17), though with pressure potentially generated by muscular action:

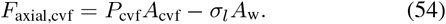

Substituting relationships for pressure (Eqn. 53), area (Eqns 42 and 45), and longitudinal stress (Eqn. 51) gives:

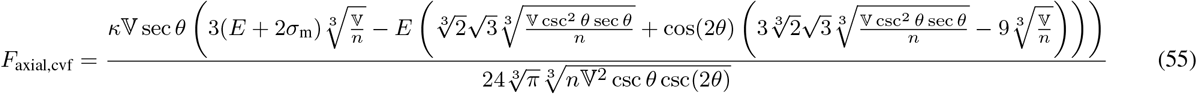

**Fig. 6.**
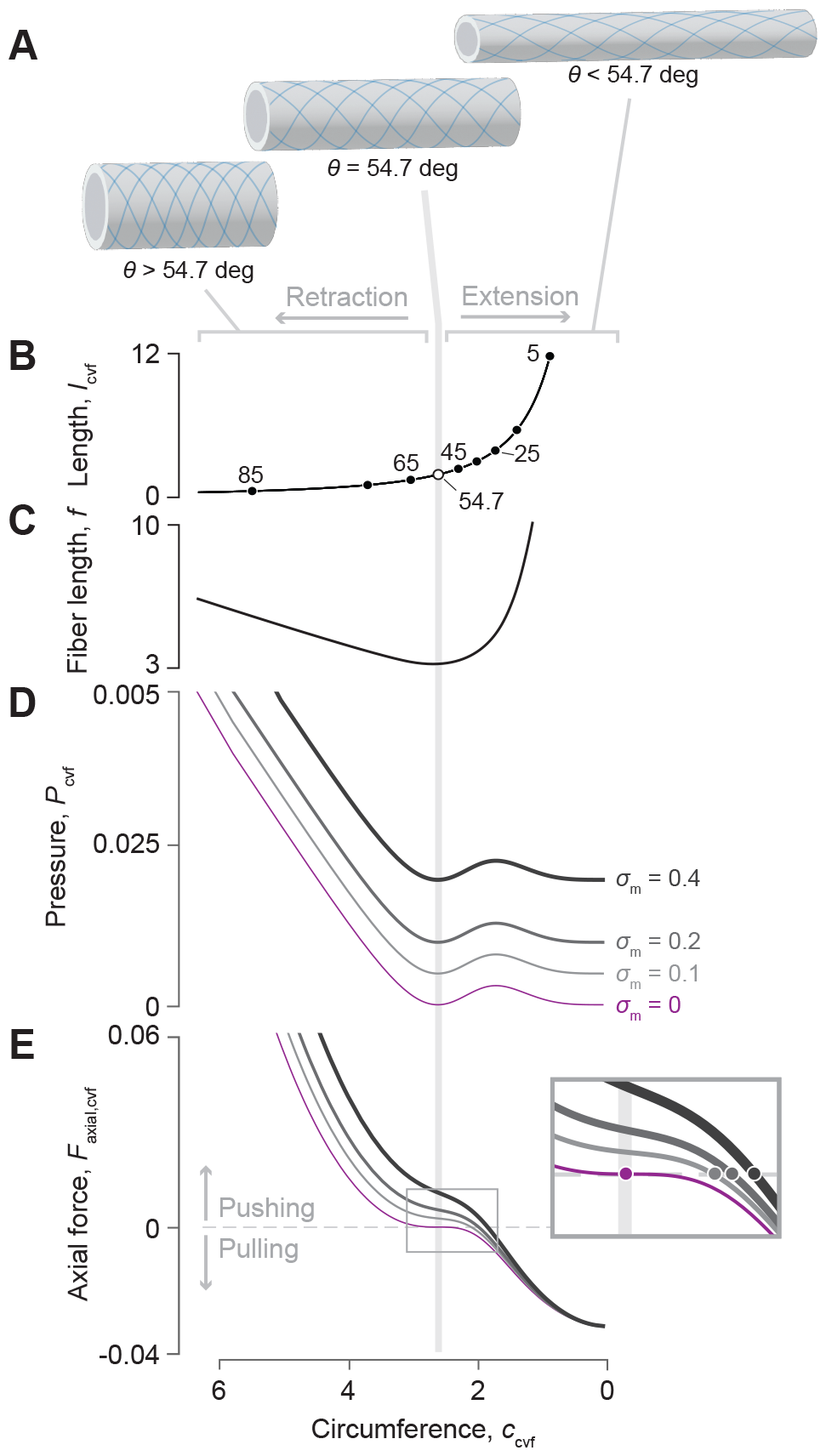
Shape, pressure, and axial force for a cylindrical hydrostat with extensible cross-helical fibers. (A) A schematic drawing illustrates changes in shape that accompany differences in helix angle. (B–E) Parametric plots of body length, fiber length, pressure and axial force with respect to circumference, are all functions of fiber angle using parameter values (*n* = 1, 𝕍 = 1, *κ* = 0.1 and *E* = 1). (B) Body length, shown with helix angle, at intervals of 10 deg (filled circles) and at 54.7 deg (open circle). The (C) fiber length (Eqn. 38) and (D) pressure (Eqn. 53) show minimum values where *θ* = 54.7 deg, with pressure increasing with muscular stress (*σ*_m_, thicker curves for greater tension with the case of no tension highlighted in violet). (E) The axial force (Eqn. 55) depends on the stress generated by circular muscles. At zero force (dashed line), the force generated by stress within the fibers balances the muscular forces, which is achieved at smaller values of circumference (filled circles in inset) with greater muscle tension.

Plotting the predictions of axial force illustrates some of the salient features of a fiber-wound hydrostat. The passive properties of this system may be considered for the case where the circular muscles generate no tension (violet curve in Fig. 6B–E). The body generates no axial force when helix angle is 54.7 deg. Despite the Hookean elasticity of the fibers, the helical geometry generates a non-linear relationship between circumference and force that elevates the restorative forces at the extremes. For example, body shortening expands the circumference and elevates fiber tension in a manner that generates a pushing force by the body, which will work to elongate the body back to its resting form like a non-linear spring.

The action of circular muscles serves to modulate body mechanics. Contraction of circular muscles generates an axial pushing force that is higher at a given fiber angle than it was at that fiber angle when circular muscles were not activated (Fig. 6E). The circumference (and hence helix angle) at which zero axial force is achieved depends on the stress generated by circular muscles (inset in Fig. 6E). If the body were to be stretched beyond this zero-axial-force point, a pulling force would tend to return the body to its balanced state. This restorative force is principally due to the passive mechanics of the fibers as they become more nearly aligned with the axial direction, and hence become more independent of the level of circular muscle activation.

At all muscle stresses, a pressure minimum occurs at 54.7 deg, which is sometimes called the “magic angle” (Horgan and Murphy, 2018; 2022); this the angle where loads in the fibers balance the hoop and longitudinal tension in the cylinder due to pressure loads. Hoses are typically manufactured with helical reinforcing fibers at this angle to prevent sudden elongation or shortening when the hose is pressurized. It can be shown that 54.7 deg is the pressurization magic angle for axial extension under the assumptions of a thin membrane and inextensible fibers (Demirkoparan and Pence, 2015). Our equations show that in constant volume cylindrical hydrostats with extensible fibers, circular muscle stresses can carry some of the pressure load that would otherwise be carried by the helical fibers and that this load sharing causes the magic angle to move to lower angles corresponding to longer, thinner cylinders.

#### Mechanical advantage and displacement advantage on axial extension

Determining the mechanical advantage when the body extends requires a model for the input force. This force is generated by the circular muscles and is equal to the product of muscle stress and wall thickness and length of the cylindrical body (*F*_in,cvf_ = *l*_cvf_*tσ*_m_). Writing the length of the body as a function of the helix angle (Eqn. 40) yields:

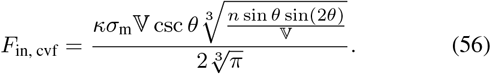

The mechanical advantage is the ratio of axial force (Eqn. 55) to input force (Eqn. 56) for the extension by circular muscles:

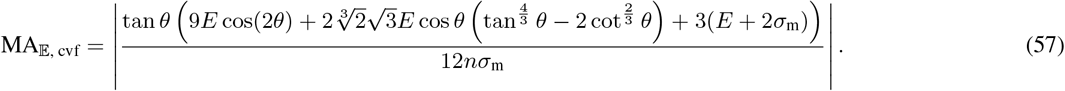

The DA on extension is a function of radius and length of the cylinder with respect to fiber angle (Eqns 40 and 41):

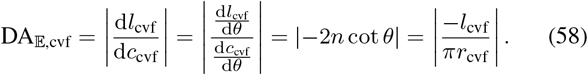

The displacement advantage here is equivalent to the cylindrical hydrostat without fibers (Eqn. 32). This is perhaps unsurprising given that a cylinder possesses only two variables, one changes through muscular action and the other is dictated by the equation for the volume of a cylinder (Eqns 40 and 41). Note that the DA here is not equal to the inverse of the MA (Eqn. 57), which reflects the storage of elastic energy in the helical fibers.

The transmission efficiency is given (using Eqns 57 and 58) by:

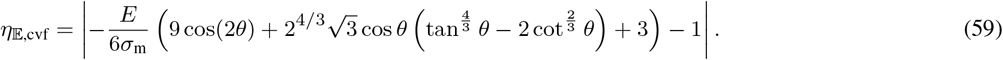

#### Mechanical and displacement advantage on axial retraction

Consider an output force in the radial direction, such as might be applied to the burrow wall of an earthworm. This is distinct from the input force on extension, which we assumed to be generated by circular muscles (Eqn. 56). Thus, the mechanical advantage on retraction is the inverse of that for extension, times a geometrical factor (2*π*) that converts circumferential to radial forces:

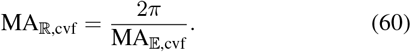

This relationship between the mechanical advantage in the two modes of operation for the worm can be seen graphically (Fig. 7D), where the mechanical advantage in retraction at 54.7 deg is 2*π* times higher than that of extension.

**Fig. 7.**
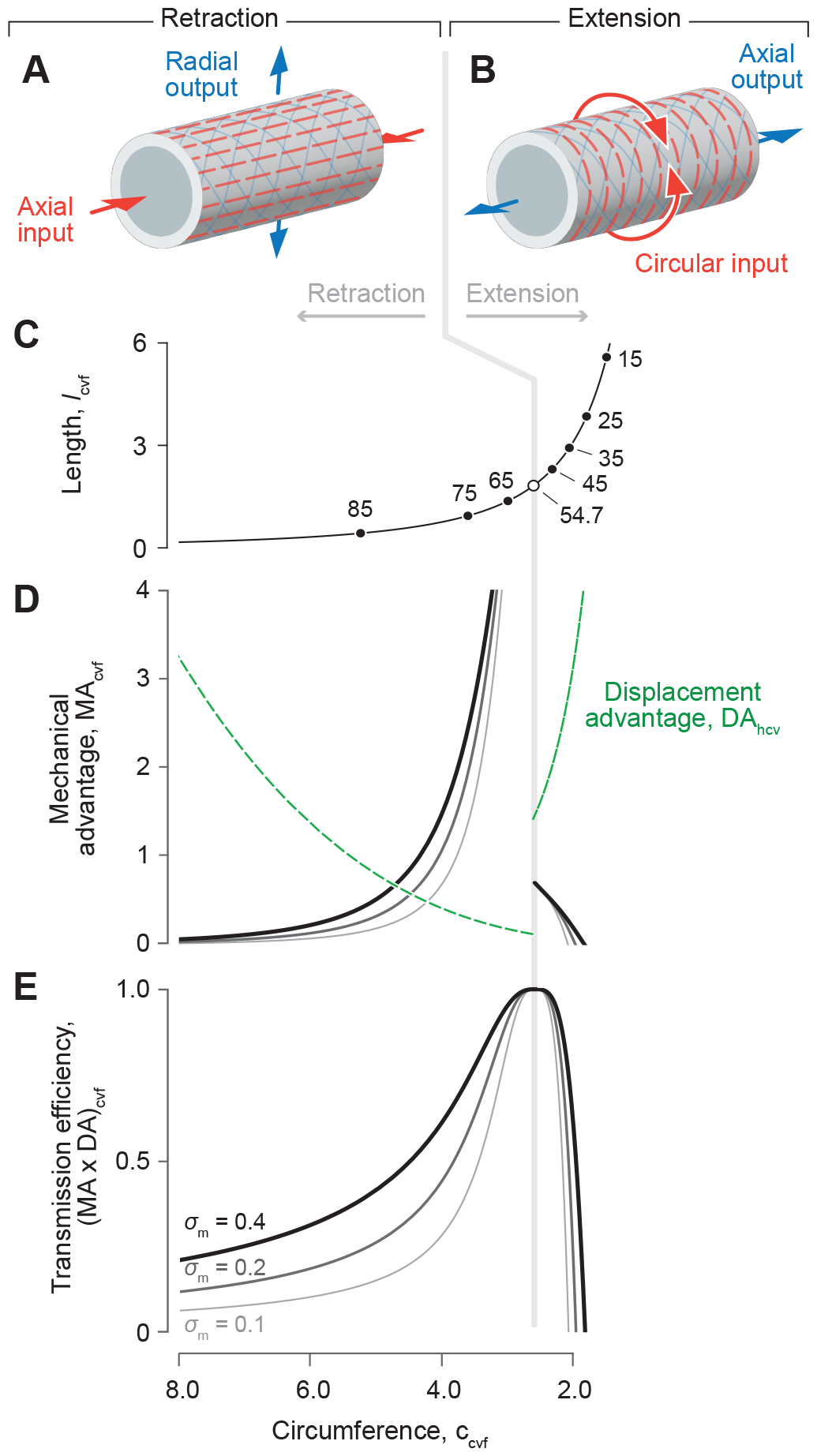
Mechanical and displacement advantage of a cylindrical hydrostat with helical fibers. (A) The body during retraction shortens (red arrows) and expands its radius (gray arrows) through contraction of longitudinal muscles (dashed red lines). (B) Circular muscles (dashed red curves) reduce the circumference (red arrows) to extend the body (gray arrows). (C–E) These changes in body shape affect a number of geometric and mechanical properties, which are expressed by parametric functions of fiber angle, plotted with respect to a reduction in circumference (towards the right) using parameter values (*n* = 1, 𝕍 = 1, and *E* = 1). (C) Body length and helix angle (intervals of 5 deg, filled circles, and at 54.7 deg, open circle) depend on circumference (Eqns 41 and 40). (D) Differences in circumference also change the mechanical advantage (gray and black, Eqns 57 and 60) and displacement advantage (green dashed, Eqns 58 and 61). Mechanical advantage additionally depends on the stress (*σ*_m_) generated by circular muscles (on extension) and longitudinal muscles (on retraction). These changes are reflected in (E) transmission efficiency, which also differs between extension (Eqn. 59) and retraction (Eqn. 62).

The displacement advantage for retraction may be derived similar to the fiberless model (Eqn. 36), but incorporates how the radius and length of the cylinder change with helix angle (Eqns 40 and 41):

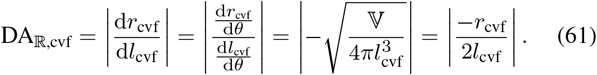

As for extension, the displacement advantage here is the same as the cylindrical hydrostat without fibers (Eqn. 36). As for the fiberless model, the displacement advantage for retraction is proportional to the inverse of that for extension [DA_ℝ,cvf_ = 1*/*(2*π*DA_𝔼,cvf_)]. Given this relationship and Eqn. 60, the transmission efficiency for retraction is:

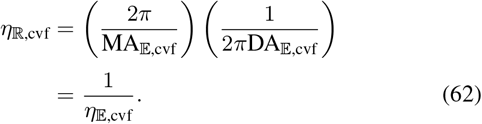

Therefore, the transmission efficiency for retraction is simply the inverse that for extension (Eqn. 59).

#### Patterns of mechanical advantage and transmission efficiency in cylindrical hydrostats with cross-helical fibers

Consider a given cylinder (constant *n, E*) with a specified *σ*_m_. As this cylinder changes shape, concomitant to changes in fiber angle, the mechanical and displacement advantage in retraction and extension (Eqns. 57-62) can be plotted (Fig. 7). Generally, displacement advantage decreases and mechanical advantage and transmission efficiency increase as 54.7 deg is approached. Transmission efficiency is equal to unity only when the helix angle is 54.7 deg, where no energy is stored in the helical fibers. There are many cases where this may be valid because a variety of biological systems maintain a fiber winding close to 54.7 deg.

However, both extension and retraction of the body stretch the helical fibers, store elastic energy, and thereby reduce transmission efficiency as the body deforms (Fig. 7E). Factors affecting mechanical advantage can be identified from Eqns. 57 or 60. Energy storage will be greater further from 54.7 deg and is affected by both circular muscle stress and the stiffness of the helical fibers. Specifically, the mechanical advantage curve will be less than the inverse of displacement advantage to a greater extent when the fiber stiffness creates resistance that is substantially greater than the force of the circular muscles.

#### Mechanical advantage at 54.7 deg

The mechanical advantage at 54.7 deg is much greater on retraction than extension in the example plotted in Fig. 7, but in general this will not be the case. Specifically, using Eqns. 43, 57, and 60, we can relate mechanical advantage to the number of windings *n* and to the length to radius ratio:

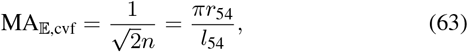

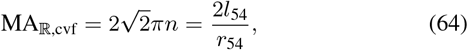

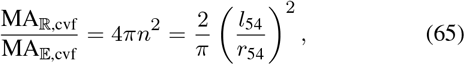

where *l*_54_ and *r*_54_ are the length and radius of a constant volume, cylindrical, fiber wound hydrostat when fiber angle is 54.7 deg. Note that at 54.7 deg, the mechanical advantage formulas for constant volume fiber wound hydrostats are identical to those for constant volume fiberless hydrostats (compare with Fig. 5 and Eqns 35, 37). Furthermore, the ratio of mechanical advantage in retraction compared to extension at 54.7 deg, i.e. at a given shape, depends on *n*^2^ (Eqn. 65). This means that mechanical advantage in retraction will always be larger than in extension when the cylinder at 54.7 deg has 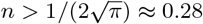. This number of fiber wrappings per length occurs at a length to radius ratio, solving from Eqn. 43, of 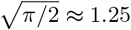. Longer, thinner cylinders at 54.7 deg will have higher mechanical advantage in retraction, i.e. generate higher radial output forces relative to axial input forces; and shorter, wider cylinders will have a higher mechanical advantage in extension, i.e. generate higher axial output forces than input circular forces. The example plotted in Fig. 7 has *n* = 1 and thus has a length to radius ratio of 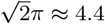, calculated from Eqn. 43. It will be interesting to compare length to radius ratios of biological cylinders operating close to 54.7 deg to see whether the length to radius ratio confers a higher mechanical advantage in retraction or extension. It will also be interesting to compare the predictions about mechanical advantage from the theory with and without helical fibers, a comparison we will undertake in the discussion.

#### Comparing mechanical advantage in a fiber wound constant volume hydrostat with a fiberless constant volume hydrostat

To compare the predictions of these two hydrostat models we calculate mechanical advantage in extension and retraction using Eqns. 43, 57, and 60, but with the assumptions that the fibers have no stiffness thus making it impossible for them to affect the mechanics by storing energy. The mechanical advantages when *E* = 0 are:

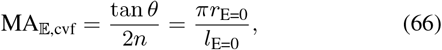

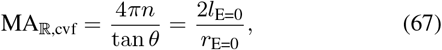

where *l*_E=0_ and *r*_E=0_ are the length and radius of a constant volume fiber wound cylindrical hydrostat with fibers with zero or low stiffness. We can see that when the fibers are not stiff, the mechanical advantages for fiber wound constant volume hydrostats are the same as those predicted for fiberless constant volume hydrostats in Fig. 5 (see also Eqns 35 and 37). To further estimate the effects of muscle stress and fiber stiffness one can plot an overlay of predictions from fiber wound and fiberless cases. First, plot MA_𝔼,cvf_ and MA_ℝ,cvf_ from Eqns. 57 and 60 against the length to radius ratio from Eqn. 43, all of which are parametric functions of *θ*. Then overlay a plot of MA_𝔼,cv_ and MA_ℝ,cv_ from Eqns 35 and 37 against the length to radius ratio. On such plots one can vary fiber stiffness and fiber stress to discover that the higher the muscle stress relative to the fiber stiffness, the closer are the predictions of mechanical advantage in the fiber wound and fiberless cases. One can also see that higher fiber stiffness generates lower mechanical advantage at a given length to diameter ratio. We leave this plot as an exercise for the reader.

### DISCUSSION

Our mathematical models consider the transmission of force and displacement by a variety of hydrostatic skeletons. The principle aims for this modeling were to examine how variable gearing emerges among hydrostatic skeletons and to evaluate the conditions where the mechanical advantage is equivalent to the inverse of the displacement advantage (Table 1). A prevailing theme in the predictions of these models is that hydrostatic skeletons transmit force with variable gearing as the structure deforms (Figs. 2H, 3C, 4B–C, 5C–F, and 7D). Our modeling also reveals the importance of identifying input force and displacement axes and the contribution of energy storage and dissipation to this variable gearing. Incorporating these considerations into calculations of mechanical advantage, displacement advantage, and transmission efficiency inform our understanding of biological soft skeletons and the design of engineered devices. While the present results offer an analytical framework, the utility of our models remains to be tested experimentally.

### Variable gearing

We found that variable gearing emerges in hydrostatic skeletons in different forms. The hydraulic press exhibits no variable gearing: the rigid walls of its two chambers yield constant mechanical and displacement advantage (Fig. 2A–C, Eqns 9–10). The hydraulic press consequently operates like a rigid joint (Fig. 1F), despite transmitting pressure through a liquid medium. However, replacing one of these chambers with the deformable McKibben actuator introduces variable gearing as a function of the helical winding (Eqns 12, 21 and Fig. 2D–H). The rapid, vertically asymptotic increase in the displacement advantage of the McKibben (Fig. 2H) may appear superficially similar to that of a cylindrical hydrostat (Fig. 5), as previously proposed (Kier and Smith, 1985), but is different because it occurs towards the magic angle, in the middle of the range of possible cylinder shapes. The special condition in the middle of the shape range is previously unrecognized in the biological literature and applies to all helically wrapped pressurized structures considered (Figs. 5 and 7).

The tube foot offers perhaps the most intriguing form of variable gearing. The combination of the ampullar and fiber winding geometries of the stem result in a maximum mechanical advantage in the middle of stem extension (Fig. 3C). This is unlike the monotonic decline in mechanical advantage of the McKibben actuator (Fig. 2H) and cylindrical hydrostat (Fig. 5C). Thus, the relative dimensions of the ampulla and tube feet could potentially serve to maximize force transmission at a length that corresponds to the power strokes that drive the bouncing gait of seastars (Ellers et al., 2014; Heydari et al., 2020; Ellers et al., 2021). Additionally, perhaps the stem length that maximizes force transmission corresponds to the maximum force in the length-tension relationship for the muscles in the wall of the ampulla. If so, the output force could be maximized through a combination of muscle tension and skeletal force transmission.

Another previously unrecognized phenomenon is that gearing depends on the direction of muscular input. For a constant volume circular hydrostat, we define the input displacement by circular muscles as a change in circumference, which contrasts with the previous convention of modeling changes in radius or diameter (Quillin, 1999; 2000; Kier, 2012; Law et al., 2014; Kurth and Kier, 2014; 2015). This distinction changes the anticipated displacement and mechanical advantage. For example, the displacement advantage of a circular muscle in a fixed-volume extending cylindrical hydrostat is less than that of a transverse muscle by a factor of *π*. Consequently, transverse muscles in squid tentacles contribute to speedy extension about 3 times more than would circular muscles.

Insight can also be gained into earthworm mechanical behavior by comparing mechanical advantage in extension and retraction for earthworm segments. From formulas for fiberless constant volume hydrostats in Fig. 5 (Eqns 35 and 37):

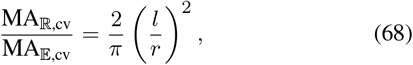

By plugging in 0.27, calculated from Quillin (1998), for *l/r* of an earthworm segment, in Eqn. 68 we infer that the mechanical advantage in retraction in earthworms is just 0.046 times the mechanical advantage in extension. This direct comparison between mechanical advantage in retraction vs. extension has not previously been made (Quillin, 1998; Kurth and Kier, 2014; 2015). We have used a calculation for constant volume fiberless hydrostats to make this inference, however, these earthworms are fiber wound. But, at 54.7 deg, we showed that the ratio of mechanical advantages predicted for fiber wound and fiberless constant volume hydrostats are the same (compare with Eqn. 65). This equivalence at 54.7 deg should not be too surprising because we know that at 54.7 deg the fiber wound cylinder is at equilibrium and the fibers are not stretched and not storing energy. Thus, for angles close to 54.7 deg we can infer that earthworm segment dimensions generate more mechanical advantage in extension than in retraction. One intriguing possibility is that earthworms can vary the mechanical advantage on extension and retraction by cooperating among segments. To the degree that segments may behave as longer coordinated groups they might be able to adjust the effective length to gain mechanical advantage in extension, but this proposition requires experimental testing.

Our fiber-wound models show that the helical fibers create a reference point at 54.7 deg relative to which predictions about mechanical and displacement advantage can be made. For instance, our model of a constant volume helically wrapped hydrostat shows that for a specific set of parameters the mechanical advantage is higher when retraction generates radial output forces just above 54.7 deg than when extension generates axial output forces just below 54.7 deg (Fig. 7D). The relative size of mechanical advantage close to 54.7 deg is however set by the ratio of length and radius (Eqn. 65). Our model has reset the perspective regarding MA and DA from one focused primarily on the length to diameter ratio to one focused on the fiber angle and its interaction with muscular forces. Just as in the McKibben, in which the structure can only be understood with respect to the fiber angle, and because the McKibben operates in two modes - extending or retracting towards the 54.7 deg length, a constant volume cylindrical hydrostat’s mechanical behavior must be analyzed relative to the fiber angle. Further models with time sequenced applications of longitudinal and circumferential muscle forces, muscle properties and assumed external forces must be developed to fully utilize our worm model.

### The relationship between mechanical advantage and displacement advantage

Energy relates mechanical and displacement advantage. If output work (*l*_out_*F*_out_) generated by a skeleton is equal to its input work (*l*_in_*F*_in_), then MA and DA are inverses which corresponds to an ideal transmission efficiency (Eqn. 3). Therefore, geometric modelling offers a comprehensive understanding of the transmission of mechanical work through a hydrostatic skeleton when energy dissipation or storage is negligible. This insight applies also to rigid skeletons.

There are two ways that a skeleton may fail to transmit all input work to the system’s output. First, energy may be dissipated as the structure deforms. For example, tube foot inflation requires fluid flow through small channels that may incur viscous losses. Similar dissipation from hydrodynamic drag has been modeled as a major factor in the force transmission of the raptorial appendage of stomatopods, due to its rapid motion (McHenry et al., 2012; 2016). Second, some of the input energy may be stored elastically. We have considered this through examples of the extension of the longitudinal muscles in the stem of the tube foot (Fig. 4) and the extension of helical fibers in the body wall of a worm (Fig. 7). The degree to which such storage occurs must be determined with experiments. For example, a preparation where the lumen of the ampulla is pressurized to a controlled extent and the muscles are rendered inactive would provide the opportunity to control the input force within a tube foot. Measurements of the output force and geometry of the tube foot would allow for measurements of the mechanical advantage, which could be compared with the predictions of our model to evaluate the extent of energy storage and dissipation in the tube foot system.

Our consideration of energetics both compliments and contrasts with previous attempts to determine the mechanical advantage by inference from the displacement advantage (Kier and Smith, 1985; Quillin, 1999; Kurth and Kier, 2014; 2015). Kier and Smith (1985) used an analogy to lever systems when they proposed that a high displacement advantage in cylindrical fixed-volume hydrostats should yield a low mechanical advantage, an approach that was adopted in other considerations of soft skeletons (Wainwright, 1988; Quillin, 2000; Vogel, 2013). For earthworms, Kurth and Kier (2014; 2015) explicitly assumed MA as the inverse of DA, with the displacement advantage calculated from morphometrics. By modeling the mechanics of a range of hydrostatic skeletons, we have show how the mechanical advantage may calculated as the inverse of the displacement advantage in a variety of systems, but only if energy is neither stored nor dissipated by the system.

### Geometric modeling of hydrostatic skeletons

The variety of helically wrapped structures presently considered offers the opportunity to expand on the traditional presentation of geometric models. Classic work (Clark and Cowey, 1958) emphasized how the volume enclosed by a cylinder (Eqn. 7) varies with the helix angle and the graphical presentation of this relationship has served as the centerpiece of textbook introductions of soft skeletons (Wainwright, 1976; Alexander, 1983; Vogel, 2013). However, this presentation can be difficult to reconcile with a cylindrical hydrostat that has a fixed volume and extensible fibers. A fixed volume requires that the system remain at one level along the ordinate. An extension of the fibers graphically shifts the operating point towards longer or shorter cylinder when helix angle is less than or greater than 54.7 deg (blue arrows in Fig. 8A). Shortening of the fibers has the opposite effect on cylinder length, moving the cylinder length towards the 54.7 deg length. An alternate graphical presentation shows the fiber length and helix angle varying along the contours of a fixed volume (Fig. 8B). Hence, shape change, fiber length change and helix angle covary (Fig. 8C). This new representation clarifies changes in a deforming hydrostat.

**Fig. 8.**
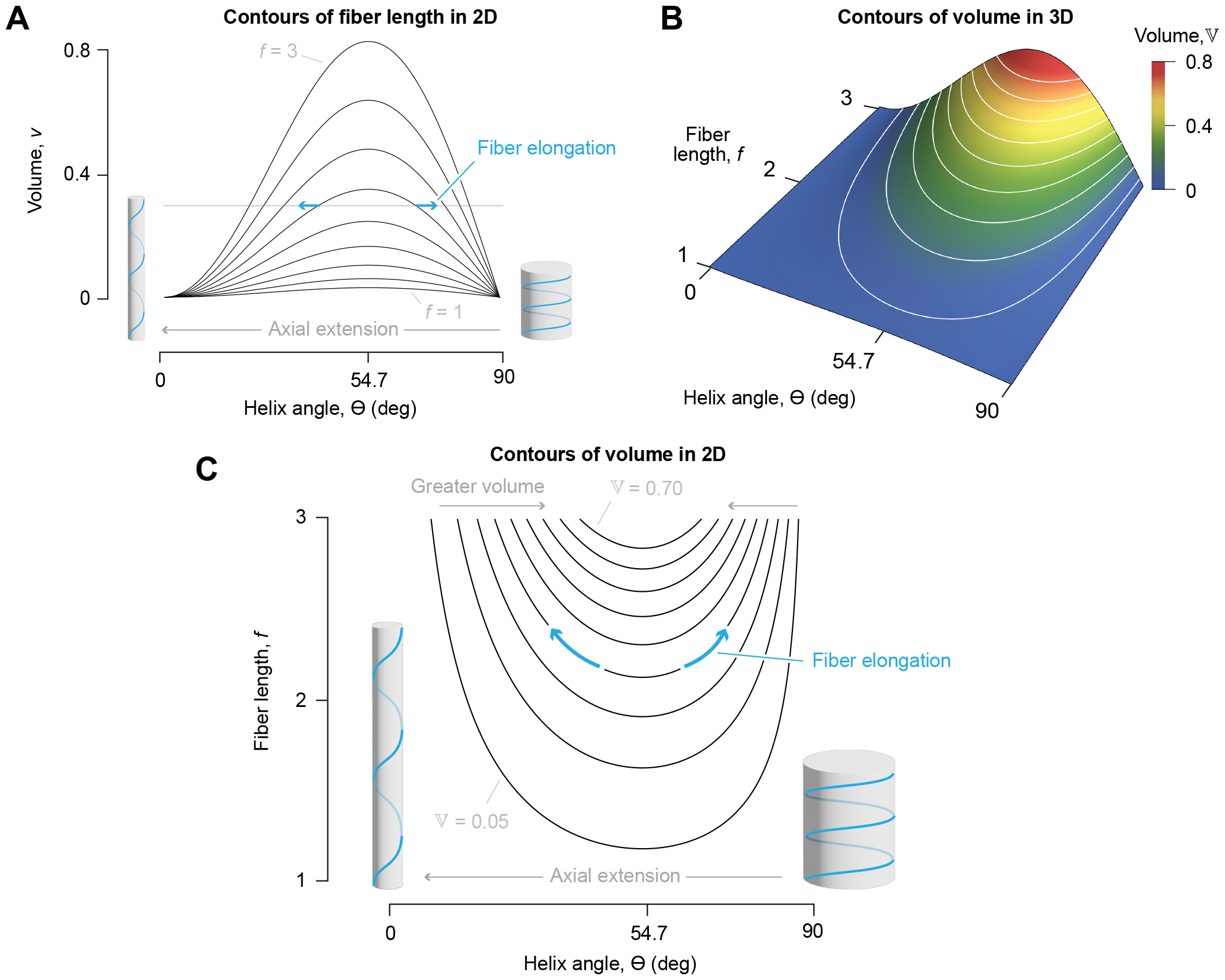
Graphical representations of the relationships between helix angle, fiber length through one helix rotation, and enclosed volume of fiber-wound cylinders (Eqn. 7). (A) Volume has traditionally been shown as a function of helix angle (Clark and Cowey, 1958), with a family of volume curves at equal intervals of fiber length (1 *≤f ≤*3). A constant volume cylinder (horizontal gray line) requires crossing fiber-length curves as the fibers elongate (e.g., along the blue arrows) to change shape. (B) The enclosed volume may alternatively be shown as a 3-dimensional plot with the helix angle in a surface plot having contours at equal intervals of volume (0.05 ≤𝕍 ≤0.70). (C) Similar contours (0.05≤ 𝕍≤ 0.80) are shown in two dimensions as curves dependent on fiber length and helix angle. In this presentation, deformation of the structure that alters helix angle away from 54.7 deg must elongate fibers (e.g., along the blue arrows).

Regardless of presentation, the present work compliments previous classic geometric models of fixed-volume cylindrical bodies (Cowey, 1952; Clark and Cowey, 1958; Kier and Smith, 1985; Kier, 2012). Our derivations similarly show that the same displacement advantage is predicted for a constant-volume hydrostat with (Eqn. 58), or without, helical fibers (Eqn. 32). In contrast, the mechanical advantage is additionally dependent on energy storage in helical fibers. Moreover, when volume is not constant, both mechanical and displacement advantage depend on the fiber angle, as illustrated by our analysis of the McKibben. Thus, our analysis considerably extends the previous understanding of these systems by explicitly modeling their mechanics.

### The mechanics of helical fiber winding

Our study deviates from prior work on soft skeletons in our explicit consideration of forces. Our models relate internal pressure to stress in helically wrapped fibers and reveals that a proportion of input work is stored as elastic energy. Such storage causes calculated MA to diverge from the inverse of DA; thus rendering potentially inoperable the widespread previous practice of using DA to estimate MA. The degree to which storage impacts force transmission will need to be measured experimentally. Our models should also enable the investigation of fiber wrapping diversity. For example, a peristalticly burrowing polychaete worm in the family Opheliidae (Law et al., 2014) has anterior circular muscles and a cuticle fiber helix angle of 45 deg, whereas an undulatory burrower has no anterior circular muscles and a cuticle fiber helix angle of 54.7 deg. Nematodes which have their own distinctive motions are pressurized, cylindrical and have reported fiber angles of 75 deg (Harris and Crofton, 1957) or 54.7 deg (Gans and Burr, 1994). Further fiber wound systems are caecilians, muscular hydrostats, fish-bodies and connective-tissue wrapped muscle. Our conceptualizations suggest a reanalysis of these systems is warranted.

It has long been recognized that fibers necessarily change length and helix angle as a constant-volume cylindrical body changes shape (Clark and Cowey, 1958). Further recognized was that fiber length is a minimum at 54.7 deg (Eqn. 5 of Kier and Smith, 1985, equivalent to our Eqn. 38 for *n* = 1). From this observation it was surmised that contracting helical muscle in a squid tentacle would cause elongating or shortening forces when the helix angle was greater or less than 54.7 deg, respectively. This corresponds to our case of zero circular muscle tension in a cylindrical hydrostat (purple line in Fig. 6D). However, we additionally modelled the patterns of axial forces when the circular muscles generate force. When circular muscles generate tension, there is a change in the critical fiber angle (and thus cylinder circumference) at which axial forces switch from pushing to pulling. Instead of occurring at 54.7 deg, the switch occurs at the fiber angle corresponding to the circumference where axial force is zero (inset of Fig. 6E). In short, axial pushing forces are generated by tension in helical fibers even at fiber angles less than 54.7 deg, when the circular muscles contract.

Muscular forces interact with the mechanics of a hydrostatic skeleton to influence the internal fluid pressure. This has been shown in experimental measurements of pressure and muscle contraction in earthworms (Seymour, 1969; Quillin, 1998; 2000). Circular muscle contraction caused pressure increases during elongation and the subsequent longitudinal contraction caused pressure decreases during shortening. This is consistent with the prediction of pressure in the region just to the right of 54.7 deg in Fig. 6D. One of the interesting consequences of the helical fibers generating forces opposing lengthening is that when there are no external axial forces acting, the length of the worm is restricted to the length that corresponds to the force balance between the helical fibers and the pressure in the coelom, resulting in the magic angle being modulated by the hoop muscle stress. This can be seen on Fig. 6E, where the axial force curves cross zero. The length of a helically wrapped worm is constrained by its helical winding.

### Biological relevance and testable predictions

Squid strike prey through the rapid extension of their tentacles. These hydrostatic structures accelerate and decelerate within a few milliseconds, but are also capable of a long extension (70%, Kier, 1982; Kier and Van Leeuwen, 1997). The initial dimensions of the tentacles (20:1 length:diameter ratio, Kier and Smith, 1985) should favor a relatively modest mechanical advantage (∼0.08, for *θ* = 54.7 deg) that declines over the course of a strike (Fig. 5C). However, these dimensions are appropriate for a high displacement advantage, which will tend to increase during the strike, provided a negligible role of fiber winding or any other connective tissue within the tentacle. If the helical fibers possess substantial stiffness, then the fiber angle changes should be quite high and thereby store elastic energy as the tentacles nearly double in length. Alternatively, fiber and muscle angles could be arranged to store energy like a spring before the strike or they may be highly compliant and thereby play little role in the tentacle’s dynamics.

Determining the role of connective tissue in the mechanics of a hydrostatic skeleton may be investigated experimentally. For example, for squid tentacles, our models suggest different modes of functioning that may be differentiated by measurements of internal pressure and electromyography. Our results suggest that preloading of helical fibers or relatively compliant fibers is necessary for a pressurized cylinder to deform to the extent exhibited by the tentacles. A forward dynamics model (Van Leeuwen and Kier, 1997) suggests that the transverse muscle arrangement with short sarcomeres is sufficient to explain the rapid strike, with a negligible role from helical fibers. Indeed, our calculations also suggest that the use of transverse rather than circular muscle increases the displacement advantage by a factor of *π*. The ultrastructure of the tentacles largely explain how the transverse muscles can achieve a maximum strain rate that is an order of magnitude faster than that of the transverse muscles of the squid arms (Kier and Curtin, 2002), but it remains unclear whether fiber winding or tendons store a significant level of elastic energy over the course of a strike.

Our modeling of echinoderm tube feet reveals a number of mechanical principles. We found that the mechanical advantage declines to zero as the helix angle approaches 54.7 deg (Fig. 3C) and that the stem can only extend further if pulled by an external force. At higher values of the helix angle, a maximum mechanical advantage occurs at an intermediate length, which presents intriguing possibilities for maximizing force output. The transmission efficiency declines when a longitudinal muscle spring is incorporated into our model because of elastic energy storage (Fig. 4D), and a high stiffness of this spring lowers the mechanical advantage and reduces the maximum elongation possible. However, the existence of the spring could aid in the control of extension by resisting the rapid increase in the displacement advantage near *θ* ∼ 54 deg. These results are rich with possibilities for experimental tests of the system that could monitor or manipulate internal pressure while measuring the deformation of these structures.

The geometry of worm bodies determines whether they are functioning with high displacement advantage or high mechanical advantage when engaged in common behaviors such as extension and retraction. Several experimental studies have documented pressures and forces involved but have not incorporated consideration of the transmission and likely storage of mechanical energy in the helical fibers despite having documented their presence. Redesign of those experiments to understand the role of the helical fibers is warranted. In particular, our equations make specific predictions of how mechanical advantage will be affected as helix angle changes. Measuring helical fiber mechanical properties and tracking helix angle and pressure during force experiments similar to those of Quillin (2000) would test whether the fibers play a role in the transmission of force. Alternatively the fibers may play a negligible role if the angle stays close to 54.7 deg or if the fibers are compliant.

Our insights into the force-transmission properties of soft skeletons open opportunities for comparisons among animals and between natural and robotic devices. In a manner analogous to the study of the functional anatomy of vertebrates (Smith and Savage, 1956; Alexander, 1983), our definitions for mechanical advantage may be applied to functional morphology of soft structures in invertebrates. Our definitions also allow mechanical comparisons among hydrostatic skeletal types and between soft and jointed skeletons. Our concepts may also aid in the design of soft robotic devices to optimize force and displacement transmission for a variety of applications.

## Acknowledgements

This paper was largely inspired by the work of W.M. Kier and benefited from conversations with him and C. Rahn. Three anonymous reviewers provided highly valuable feedback on the initial version of this manuscript.

## Contribution

This study emerged from conversations among all authors. The mathematical modelling was primarily developed by OE and MJM, with assistance from KIE and SH. The manuscript and electronic supplemental materials were written by MJM, OE, and ASJ, with critical feedback from all authors.

